# Tracking the Movement of Electronic Cigarette Flavor Chemicals and Nicotine from Refill Fluids to Aerosol, Lungs and Exhale

**DOI:** 10.1101/2020.11.13.382309

**Authors:** Careen Khachatoorian, Kevin J. McWhirter, Wentai Luo, James F. Pankow, Prue Talbot

**Author notes:** **Address correspondence to** Prue Talbot, Department of Molecular, Cell and Systems Biology, University of California, Riverside, 900 University Dr. Riverside, CA 92521, USA. **E-mail:**.

## Abstract

Electronic cigarettes (ECs) have been linked to lung diseases, including COVID-19, with little understanding of exposure, retention, and exhalation of EC aerosol chemicals. Here, flavor chemicals and nicotine were quantified in two refill fluids, their transfer efficiency to EC aerosols was determined, exhalation by human participants was measured, and chemical retention was modeled. Nicotine transferred well to aerosols irrespective of topography; however, transfer efficiencies of flavor chemicals depended on the chemical, puff volume, puff duration, pump head, and EC power. Participants could be classified as “mouth inhalers” or “lung inhalers” based on their retention and exhale of flavor chemicals and nicotine. Only mouth inhalers exhaled sufficient concentrations of flavor chemicals and nicotine to contribute to chemical deposition on environmental surfaces. These data help distinguish two types of EC users, add to our knowledge of chemical exposure during vaping, and provide information useful in treating EC-related diseases and regulating EC use.

## 1. INTRODUCTION

Electronic cigarettes (ECs) are battery powered nicotine delivery devices that produce an inhalable aerosol. The battery heats a metal coil(s) surrounded by a cotton wick saturated with fluid. The user then inhales aerosol usually containing nicotine, propylene glycol (PG), glycerol (G), flavor chemicals, metals, particulate matter, and volatile organic chemicals (VOCs) (Goniewicz et al., 2013; Goniewicz et al., 2014; Trehy et al., 2011; Vansickel et al., 2018; Lerner et al., 2015; Pellegrino et al., 2012; Williams et al., 2013). The VOCs include toxic aldehydes, such as formaldehyde and acrolein, that are produced by thermal dehydration of glycerin and/or glycols (McAuley et al., 2012; Uchiyama et al., 2013). Many EC devices are customizable and allow the user to vary the voltage, wattage, and amperage (Bitzer et al., 2017), which can alter the transfer of fluid chemicals to the aerosol (Zhao et al., 2016) and may also increase the production of toxic reaction products (Logue et al., 2017).

The possible effects of EC use on human health have been reviewed (Pisinger and Døssing, 2014; Gotts et al., 2019), and recent infodemiological data show the occurrence of health issues in EC users over the last 7 years (Hua et al., 2020). The relationship between reported health and flavor chemicals/nicotine is of interest due to their frequent use at high concentrations (Behar et al., 2018; Omaiye et al., 2019; Davis et al., 2015; Hua et al., 2019) and reported toxicity. For example, vanillin, ethyl vanillin, and ethyl maltol are often used in EC products (Khlystov and Samburova, 2018; Tierney et al., 2016) and are cytotoxic to human pulmonary fibroblasts in the MTT assay (Behar et al., 2018). Ortho-vanillin and maltol increased secretion of IL-8 from BEAS-2B cells and decreased barrier function in human bronchial epithelial cells exposed *in vitro* (Gerloff et al., 2017). Flavor chemicals in aerosolized fluids (cinnamaldehyde, vanillin, and ethyl vanillin) were toxic to CALU3 cells after five puffs and caused dose-dependent decreases in cell viability (Rowell et al., 2017). Pure menthol, when aerosolized in a cloud chamber, increased mitochondrial protein oxidation, expression of the antioxidant enzyme SOD2, and activation of NF-κB, in air-liquid interface cultures of BEAS-2B cells (Nair et al., 2020). Some EC flavor chemicals damage human lung tissue. For example, inhalation of diacetyl leads to bronchiolitis obliterans, a serious and irreversible lung disease (Allen et al., 2016). Although not directly linked to flavor chemicals/nicotine, vaping does cause e-cigarette or vaping product use-associated lung injury (EVALI) (Balmes, 2019) and has been associated with COVID-19, which has a higher probability of occurring in those who have used ECs (Wang et al., 2020).

While most research focus has been on inhalable aerosols, EC users also exhale aerosol that settles on indoor surfaces where it accumulates as <u>EC</u> <u>e</u>xhaled <u>a</u>erosol <u>r</u>esidue (ECEAR). ECEAR contains nicotine, tobacco specific nitrosamines (TSNAs), solvents, and particles (Son et al., 2020; Khachatoorian et al., 2018; Bush and Goniewicz, 2015; Khachatoorian et al., 2019; Goniewicz and Lee, 2015; Sempio et al., 2019). ECEAR chemicals increased in concentration in a vape shop over a month-long period of monitoring, and concentrations were highest in heavily used areas (Khachatoorian et al., 2019). An EC user’s living room also had residue containing nicotine and tobacco alkaloids, albeit at a lower concentration than the vape shop. ECEAR can also accumulate away from its site of origin. Nicotine, other alkaloids, and TSNAs transferred from a vape shop in a mini mall to an adjacent business where they deposited on paper and cotton towels (Khachatoorian et al., 2018). As far as we know, no studies have looked at flavor chemicals in ECEAR, even though their concentrations are high in many in refill fluids (Hua et al., 2019). The effects of ECEAR on human health are unknown, but its accumulation in indoor environments presents the opportunity for active and passive exposure.

Given the high concentrations of nicotine and flavor chemicals in EC fluids and their demonstrated toxicity, it is important to determine how efficiently they transfer to aerosols (exposure), how well they are retained by users, and if they are exhaled into the environment where they can form ECEAR. The goal of our study was to obtain a complete overview of the movement of flavor chemicals and nicotine from refill fluids into aerosols, then into users’ respiratory systems, and finally into their exhale where it could contribute to ECEAR. To do this, we quantified flavor chemicals and nicotine in two refill fluids, then determined the effects of topography on their transfer into machine-generated aerosols. Human exposures were determined by measuring the concentrations of these chemicals in exhale and modeling retention using information on transfer efficiency and exhale.

## 2. Materials and Methods

### i. Refill Fluids

“Dewberry Cream” was purchased at a local vape shop that sold products made by refill fluid manufacturers, while “Cinnamon Roll with Cinnamon Bomb” was purchased at a local vape shop that custom mixes its refill fluids. Both shops were located in Riverside County, CA. “Dewberry Cream” by Kilo was chosen because it has many flavor chemicals, including vanillin, ethyl vanillin, maltol and ethyl maltol, and a high total flavor chemical concentration (Hua et al., 2019. “Cinnamon Roll with Cinnamon Bomb”, which we refer to as “Cinnamon Roll”, was chosen because it has only one dominant flavor chemical (cinnamaldehyde) and cinnamon-flavored refill fluids can adversely affect cultured cells (Behar et al., 2014; Behar et al., 2016; Wavreil and Heggland, 2019; Clapp et al., 2019; Fetterman et al., 2018). Each refill fluid was labeled to have 6 mg of nicotine/mL and 70/30 G/PG ratio.

### ii. EC Aerosol Production and Capture

For aerosol production, we used a SMOK Alien 220W Mod (variable voltage (0.35-8V) with two high amperage flat top 18650 batteries. The mod was used with a SMOK V8 Baby-Q2 (0.4) single coil tank atomizer inside a SMOK Baby Beast tank. The smoking machine was a Cole-Parmer Masterflex L/S peristaltic pump used with a standard or high-performance pump head. When set to 40 watts, aerosols were generated at 4.3 volts, 0.4 ohms, and 9.9 amps. When set to 80 watts, aerosols were generated at 6.1 volts, 0.4 ohms, and 14.1 amps. The tank was loaded with 3 mL of refill fluid each time aerosol was produced, and the EC was primed with three puffs. The tank was washed with water and ethanol, and the V8 Baby Beast coil was replaced between each refill fluid. Puff durations were 1, 2 and 4.3 seconds; the latter is a reported average for EC consumers (Hua et al., 2013).

The standard pump head (low volume pump head) generated a flow rate of 13 mL/sec to produce puff volumes of 13 mL (1 sec), 26 mL (2 sec), and 56 mL (4.3 sec). The high-performance pump head (high volume pump head) generated a flow rate of 40 mL/sec to produce puff volumes of 40 mL (1 sec) and 80 mL (2 sec).

For flavor analysis, aerosols were collected at room temperature in two 125 mL impingers, each containing 25 mL of isopropanol (IPA). The tank was weighed before and after aerosol production to collect a mass concentration of at least 15 mg/mL for GC/MS analysis. Aerosol solutions were collected, aliquoted, and stored at −20 °C until analyzed.

### iii. Identification and Quantification of Flavor Chemicals Using GC/MS

Refill fluids, aerosols, and exhale were analyzed by GC/MS. Internal standard-based calibration procedures similar to those described elsewhere were used (Tierney et al., 2016; Omaiye et al., 2019; Brown and Cheng, 2014), and analyses for 176 flavor-related target analytes and nicotine were performed with an Agilent (Santa Clara, CA) 5975C GCMS system. The capillary column used was a Restek (Bellefonte, PA) RXI-624Sil MS (30 m long, 0.25 mm id, and 1.4 μm film thickness). For each refill fluid sample, 50 μL was dissolved in 950 μL of isopropanol (Fisher Scientific, Fair Lawn, New Jersey, USA). Prior to analysis, 20 μL of internal standard solution (2 μg/μL of 1,2,3-trichlorobenzene in isopropyl alcohol) was added into the 1 mL diluted refill samples, the aerosol and exhaled extract aliquots. 1 μL of the sample was injected into the GC/MS with a 10:1 split. The injector temperature was 235°C. The GC temperature program for all analyses was as follows: 40°C hold for 2 min; 10°C/min to 100°C; 12°C/min to 280°C and hold for 8 min at 280°C, then 10°C/min to 230°C. The MS was operated at electron ionization mode. The ion source temperature was 226°C. The scan range was from 34 to 400 amu. Each target analyte was quantitated using authentic standard material, and an internal standard (1,2,3-trichlorobenzene) normalized multipoint calibration.

### iv. Participants

Ten of eleven recruited participants (3 women and 7 men) completed the exhale portion of the study. The average age was 21 years (SD = 2.8; median = 20; range = 18-28). The ethnicity of the participants was: eight Asian, one African American, and one Caucasian. All participants self-reported no use of combustible cigarettes for the duration of the study and were told to abstain from using ECs 1 hour before the experiment. Six of the participants had used combustible cigarettes in the past. One of the six used a cigarette once a month during the study and the other five reported no current use. Two of the participants had used cigars in the past. The inclusion criteria were: (1) experienced EC users (at least 3 months), and (2) must use at least 3 mg of nicotine in their current EC. Participants were excluded if they were: (1) pregnant or breast feeding, (2) under the age of 18 or over 75 years, (3) never users of EC nicotine, or (4) experiencing any medical conditions. All participants signed informed consent before admission into the study. The project was approved by the UCR Internal review Board (IRB # HS-12-023). Participants were coded to identify puffing topography and were compensated after four sessions of vaping.

### v. EC Exhaled Aerosol Production and Capture

Participants were asked not to use any ECs or cigarettes an hour before coming to the lab. Upon arrival, participants vigorously washed their mouths and gargled for 30 sec with water. A 2 feet piece of plastic tubing with a mouthpiece was attached to two impingers connected to each other by a short piece of tubing. Each impinger contained 25 mL of IPA. The first session

(control) involved collection of 30 puffs of exhale in the impingers at 1 puff/minute without any EC use. After the last puff, the sample was collected from each impinger and stored in glass vials for chemical analysis. The next four sessions involved using the SMOK Alien with the Baby Beast tank at 40 or 80 watts for each refill fluid (“Dewberry Cream” and “Cinnamon Roll”). The tank was primed with three puffs before each use. Volunteers were asked to use the EC at 1 puff/minute at 40 or 80 watts during different sessions. The puff duration was sampled two to three times during each session. At the end of a session, IPA was collected from each impinger and used to wash residual aerosols from inside the tubing and impingers. 1mL from each impinger was then aliquoted into GC sample vials for chemical analysis. The impingers, tubing, and V8 Baby Beast tank were washed with water and 75% ethanol and left to dry for the next session. Each volunteer was given a new tube and V8 Baby coil for each refill fluid. The coil in the tank was changed between each participant and each refill fluid. The SMOK Alien box mod and tank were changed once during the study.

### vi. Calculating Transfer Efficiency, Percent Retention, and ECEAR

To determine the transfer efficiency of flavor chemicals and nicotine, aerosol fluid flavor or nicotine concentrations were divided by the refill fluid flavor or nicotine concentrations. The transfer rate was multiplied by 100 to get percent transfer efficiency.

Tank weights, which were recorded before and after each session, were subtracted to find the total weight of EC fluid consumed. Potential mass delivered was calculated by multiplying the fluid consumed by the refill fluid flavor chemical or nicotine concentration. Actual mass delivered was calculated by multiplying the transfer rate by the potential mass delivered. Total mass in the exhaled aerosol was calculated by multiplying the fluid consumed by the concentration in the exhaled sample. The percent retention was calculated by the following equation:

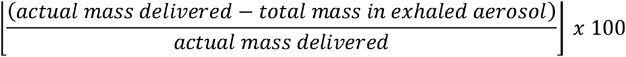

ECEAR was calculated by subtracting the percent retention from 100.

## 3. RESULTS

### i. Refill Fluid Characterization

“Dewberry Cream” and “Cinnamon Roll” had 47 and 36 flavor chemicals, respectively. Heatmaps show all flavor chemicals (y-axis) detected and quantified in the refill fluid (x-axis) (Figure 1). The total quantity of flavor chemicals and nicotine are listed at the top of each column in mg/mL.

**Figure 1:**
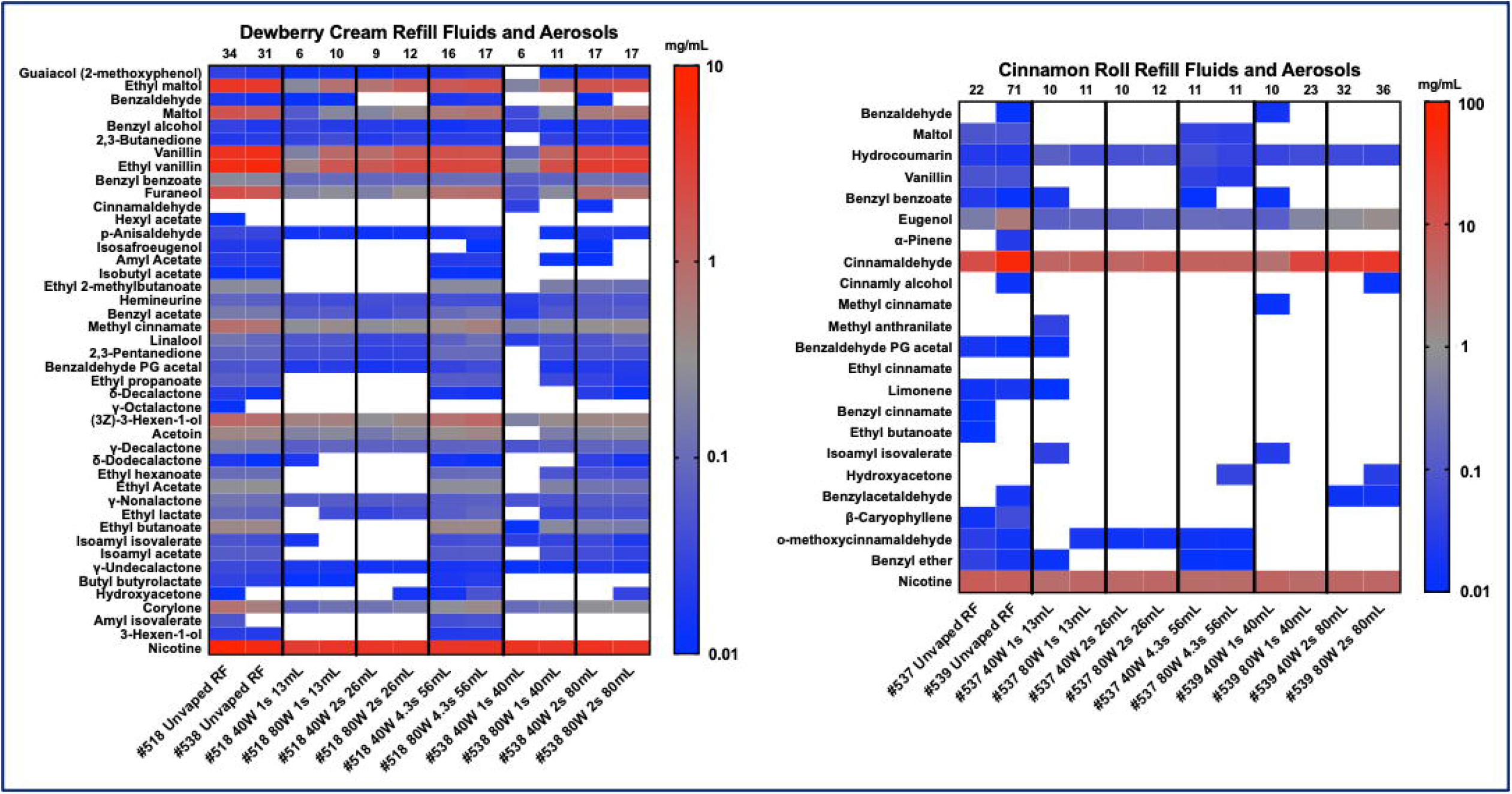
Heatmaps showing concentrations of flavor chemicals and nicotine in “Dewberry Cream” (A) and “Cinnamon Roll” (B) refill fluids and aerosols made at 40 or 80-watts. Puff durations were either 1, 2, or 4.3 seconds, while puff volume was either 13, 26, 56, 40, or 80 mL. Flavor chemicals are listed on the left y-axis and concentrations are in mg/mL. The top x-axis shows the total mg of flavor chemicals including nicotine in each column.

#### i. Dewberry Cream

“Dewberry Cream” is distributed in vape shops nationally and can be purchased online. Its flavor profile is described as mixed berries, honeydew, and cream. Bottles #518 and #538 were purchased at different times. Although their total flavor chemical concentrations varied by 3 mg, the concentrations of the dominant flavor chemicals were similar in each bottle (Fig. 1A). Dewberry Cream (#518) contained > 1 mg/mL of ethyl vanillin (6.1 mg/mL), vanillin (4.7 mg/mL), ethyl maltol (4.4 mg/mL), maltol (1.9 mg/mL), furaneol (2.1 mg/mL), and (3Z)-3-hexen-1-ol (1 mg/mL). Although labeled as 6 mg/mL of nicotine, the actual concentration was 8.7 mg/mL. “Dewberry Cream” (#518) was the refill fluid that participants used to create exhaled aerosols. Both “Dewberry Cream” #518 and #538 were used to create aerosols to determine transfer efficiency.

#### ii. Cinnamon Roll with Cinnamon Bomb

“Cinnamon Roll with Cinnamon Bomb” was custom mixed for us on two occasions at a local vape shop in Riverside, CA. The mixture’s flavor profile was described as mostly cinnamon with some sweet flavors. Bottles #537 and #539 were not identical and had different concentrations of cinnamaldehyde and eugenol, the two dominant flavor chemicals (Figure 1B). “Cinnamon Roll” (#537) contained 13.5 mg/mL of cinnamaldehyde, and although labeled as 6 mg/mL nicotine, the actual concentration was 7.5 mg/mL. Other flavor chemicals in “Cinnamon Roll” were eugenol (0.4 mg/mL), maltol (0.09 mg/mL), vanillin (0.08 mg/mL), and hydrocoumarin (0.02 mg/mL). “Cinnamon Roll” (#537) was the refill fluid that participants used to create exhaled aerosols. Both “Cinnamon Roll” #537 and #539 were used to create aerosols to determine transfer efficiency.

### ii. Aerosol Characterization

#### i. E-Liquid Aerosolized and EC Setting

The amount (mg) of refill fluid aerosolized with the Smok Alien V8 baby beast tank from 30 puffs with the low and high-volume pump heads is shown in Table 1.

**Table 1:**
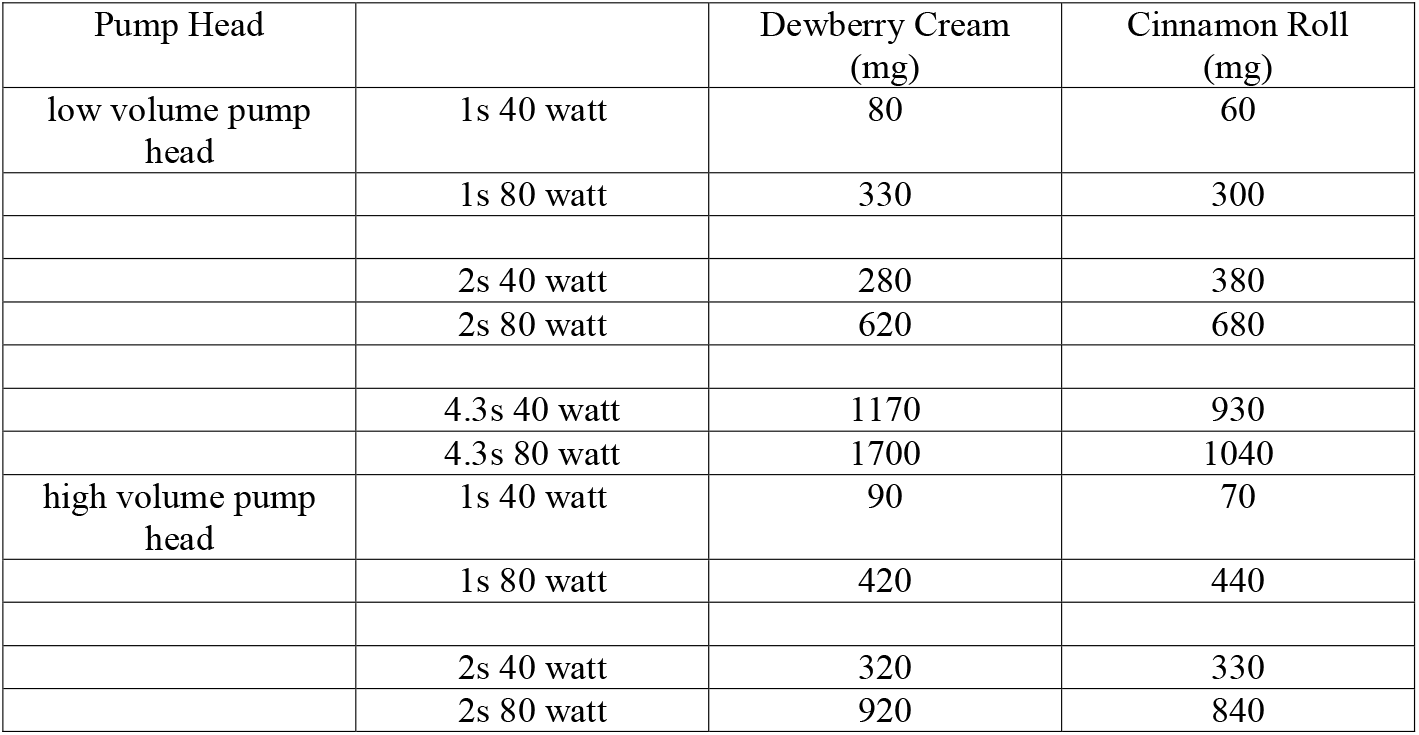
Amount of refill fluid aerosolized at different EC settings

### iii. Transfer Efficiency

Figure 2 shows that the transfer efficiency of the major flavor chemicals and nicotine from the refill fluids to the aerosol was affected by topography. Two different pump heads were used to create aerosol to get a range of flow rates. The low flow rates are shown in Figure 2A, C, E, G, I, K, M, O and Q, while the high flow rates are shown in Figure 2B, D, F, H, J, L, N, P and R. The 40-watt setting is almost always lower in transfer efficiency than the 80-watt setting.

**Figure 2:**
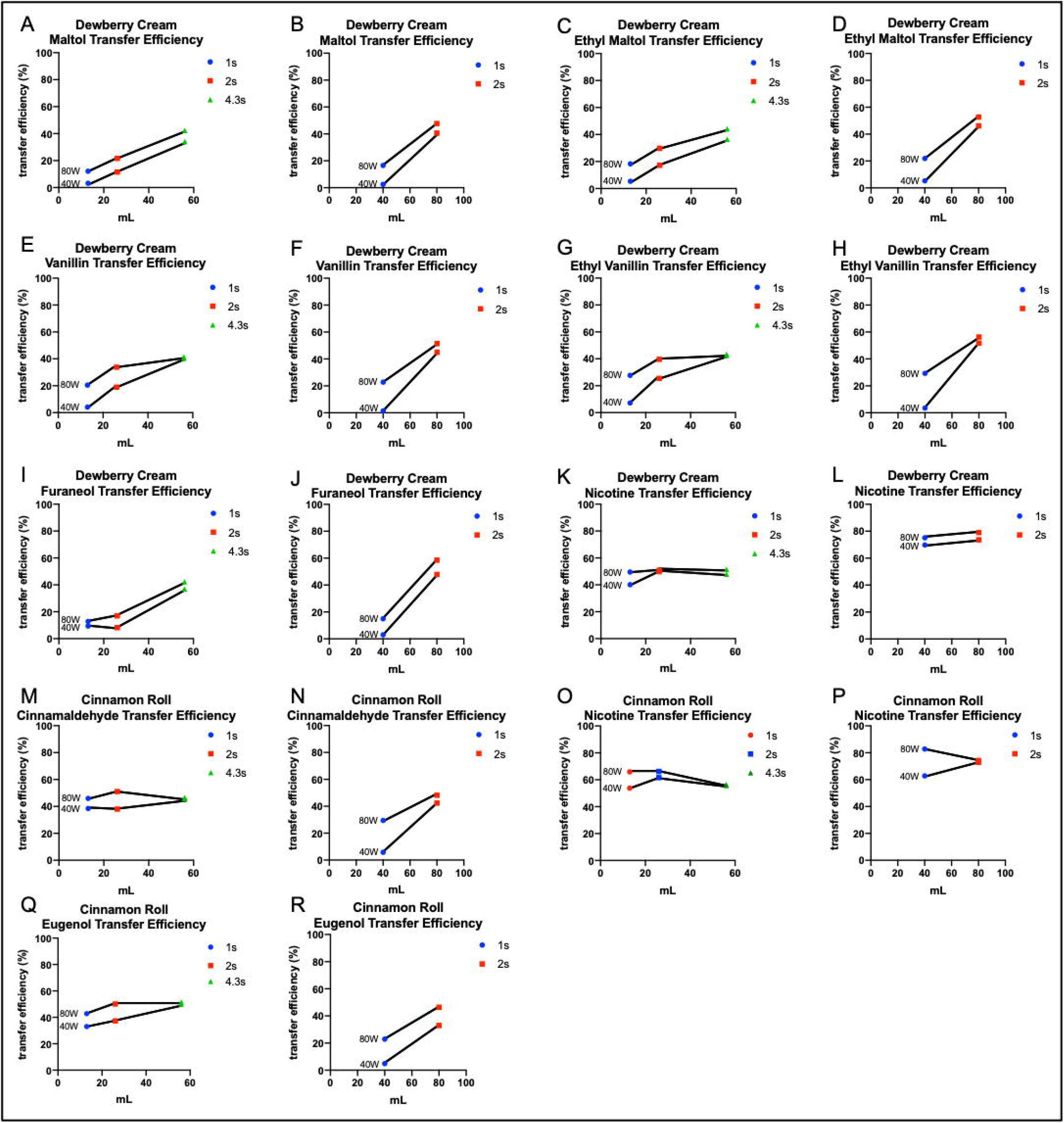
Transfer Efficiency of major flavor chemicals in “Dewberry Cream” and “Cinnamon Roll”. Aerosols made with the low volume pump head are shown in A, C, E, G, I, K, M, O and Q, while aerosols made with the high volume pump head are shown in B, D, F, H, J, L, N, P and R. Volume is shown on the x axis and transfer efficiency (in percentage) is shown on the y axis.

### iv. Low Flow Rate

Maltol, ethyl maltol, vanillin, ethyl vanillin, and furaneol have similar patterns for each puff duration, EC setting, and puff volume (Figure 2A, C, E, G, I). Lower puff durations had a lower transfer efficiency than higher puff durations with the low volume pump head. Cinnamaldehyde was consistently between 38 −51% transfer efficiency for each puff volume (Figure 2M). Similarly, eugenol was between 33-51%. Finally, nicotine was consistently between 40-51% and 54-66% for “Dewberry Cream” (Fig. 2K) and “Cinnamon Roll” (Fig. 2O), respectively.

### v. High Flow Rate

Maltol (Fig. 2B), ethyl maltol (Fig. 2D), vanillin (Fig. 2F), ethyl vanillin (Fig. 2H), and furaneol (Fig. 2J) have similar patterns for each puff duration, EC setting, and puff volume. The lower puff duration (1 second), had a lower transfer efficiency than the higher puff duration (2 seconds). When using the high flow pump head, 1 second puff durations were lower or equal to the transfer efficiency of 1 second puff durations with the low flow pump head. Cinnamaldehyde (Fig. 2N) had a 5% transfer efficiency for the 1 second 40-watt setting, but when the wattage increased to 80, the transfer efficiency increased to 30%. In a similar pattern, eugenol had a 5% transfer efficiency for the 1 second 40-watt setting, but when the wattage increased to 80, the transfer efficiency increased to 23%. Nicotine had a consistent transfer efficiency between 70-79% for “Dewberry Cream” (Fig. 2L) and between 63-82% for “Cinnamon Roll” (Fig. 2P).

### vi. Exhaled Aerosol

#### i. Puff Duration

Participants used the SMOK Alien at a low and high wattage with “Dewberry Cream” and “Cinnamon Roll” refill fluids. Control analysis of exhale (no EC use) showed minimal to no detectable levels of nicotine or flavor chemicals (Supplementary Table 1). Each participant took 30 puffs at 1 puff/minute of either “Dewberry Cream” or “Cinnamon Roll” refill fluid at 40-watts or 80-watts, and puff duration was sampled for each participant (Figure 3A and 3B). A total of 5 puffing sessions/participant, including one control session, was documented and analyzed. Most participants had puff durations between 0.5-2 s. For each participant, puff durations for both wattages were similar with deviations generally no more than 0.5 seconds.

**Figure 3:**
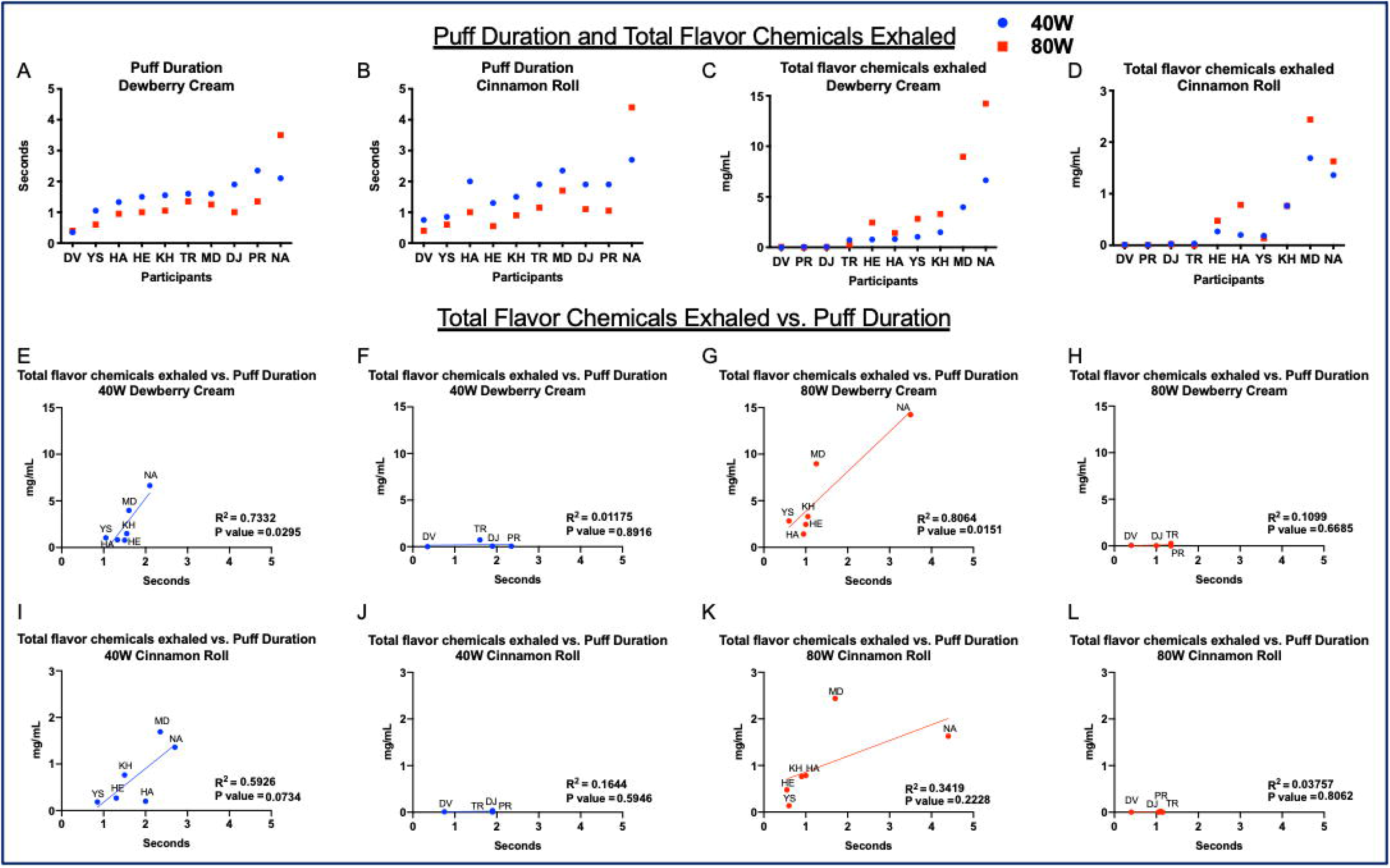
Participant topography. A and B show each participant’s puff duration for Dewberry Cream and Cinnamon Roll. C and D show the concentration of the total flavor chemicals exhaled (mg/ml). E through L show the relationship between the total flavor chemicals exhaled and puff duration. Mouth inhalers are shown in E, G, I, and K, while lung inhalers are shown in F, H, J, and L.

Puff duration was generally longer for the 40-watt setting for both refill fluid flavors. One participant, NA, a higher puff duration (2.1s for 40W “Dewberry Cream”, 3.5s for 80-watt “Dewberry Cream”, 2.7s for 40-watt “Cinnamon Roll”, and 4.4s for 80-watt “Cinnamon Roll”) than the others at both settings and for both refill fluid flavors. The average puff duration for all participants was 1.4 ± 0.27 s. The average puff duration for “Dewberry Cream” (DC) at 40-watts = 1.5 ± 0.56 s, DC at 80-watts = 1.2 ± 0.85 s, CR at 40-watts = 1.7 ± 0.62 s, and CR at 80-watts = 1.3 ± 1.15 s.

#### ii. Total Flavor Chemicals

Heatmaps of participants’ exhaled aerosol chemical concentrations are shown in Supplementary Figure 1. Total flavor chemical concentration in the exhale was generally higher for the 80-watt setting (Figure 3 C, D). Four participants (DV, PR, DJ, and TR) exhaled almost no flavor chemicals, and we categorized these as “lung inhalers” (i.e., all of the aerosol likely reached the alveoli of the lungs). Six of the participants (HE, HA, YS, KH, MD, and NA) exhaled a fraction of the flavor chemicals that they inhaled, and these were categorized as “mouth inhalers” (i.e., intake went mainly into the mouth but did not fully penetrate into the lungs). The mouth inhalers exhaled 1 to 15 mg of the total flavor chemicals. The average concentration of flavor chemicals exhaled for Dewberry Cream and Cinnamon Roll at 40-watts by mouth inhalers increased as wattage increased (Dewberry Cream = 2.5 ± 2.4 to 5.5 ± 5 mg and Cinnamon Roll = 0.7 ± 0.6 mg to 1 ± 0.8 mg). The average concentration of flavor chemicals exhaled by lung inhalers was low and similar between the two wattages (Dewberry Cream 40 watts = 0.2 ± 0.3 mg, 80-watts = 0.09 ± 0.1 mg; Cinnamon Roll = 40-watts = 0.02 ± 0.01 mg, 80-watts = 0.007 ± 0.007 mg). The average total flavor chemicals exhaled for all participants was 1.5 ± 1.87 mg and averages increased with increasing wattage (Dewberry Cream = 40-watts = 1.6 ± 2.13 mg, 80-watts = 3.4 ± 4.68 mg, and Cinnamon Roll = 40-watts = 0.5 ± 0.61 mg, 80-watts = 0.6 ± 0.83 mg).

#### iii. Total Flavor Chemicals Exhaled Vs. Puff Duration

The total flavor chemicals exhaled vs. puff duration for each refill fluid and EC setting are shown in Figure 3E-L. Participants were separated based on whether they were “mouth inhalers” (Fig. 3E, G, I, K) or “lung inhalers” (Fig. 3F, H, J, L). Dewberry Cream refill fluid puffed at 40 (Fig. 3E) and 80 watts (Fig. 3G) had significant correlation for the amount of flavor chemicals exhaled and puff duration for mouth inhalers. Cinnamon Roll puffed at the 40-watt (Fig. 3I) was not correlated flavor chemical concentration but had a p value close to 0.05. Cinnamon Roll at 80 watts (Fig. 3K) was not significant, but when reanalyzed without the outlier (MD), the p value decreased from 0.22 to 0.03 indicating a correlation. There was no correlation of exhaled chemicals and puff duration for the lung inhalers (3F, H, J, L).

#### iv. Fluid Consumed Vs. Flavor Chemicals Exhaled

The average fluid consumed for all participants was 567 ± 112 mg. The average fluid consumed was lower at the 40-watt setting and higher at the 80-watt setting (DC 40-watts = 429 ± 261 mg, DC 80-watts = 705 ± 308 mg, CR 40-watts = 424 ± 197 mg, and CR 80-watts = 713 ± 462 mg). Figure 4A-H shows the relationship between the amount of refill fluid consumed and the concentration of flavor chemicals exhaled. For lung inhalers there was no correlations between how much fluid was consumed and the amount of flavor chemicals exhaled (Fig. 4B, D, F, H). For mouth inhalers, “Dewberry Cream” at the 40-watt setting showed significant correlation (R^2^ = 0.85, p = 0.008) (Fig 4A). Mouth inhaler data appeared to be linearly related but were not significantly correlated. However, when Figure 4G was re-analyzed without the outlier (MD), the p = 0.03, indicating significance.

**Figure 4:**
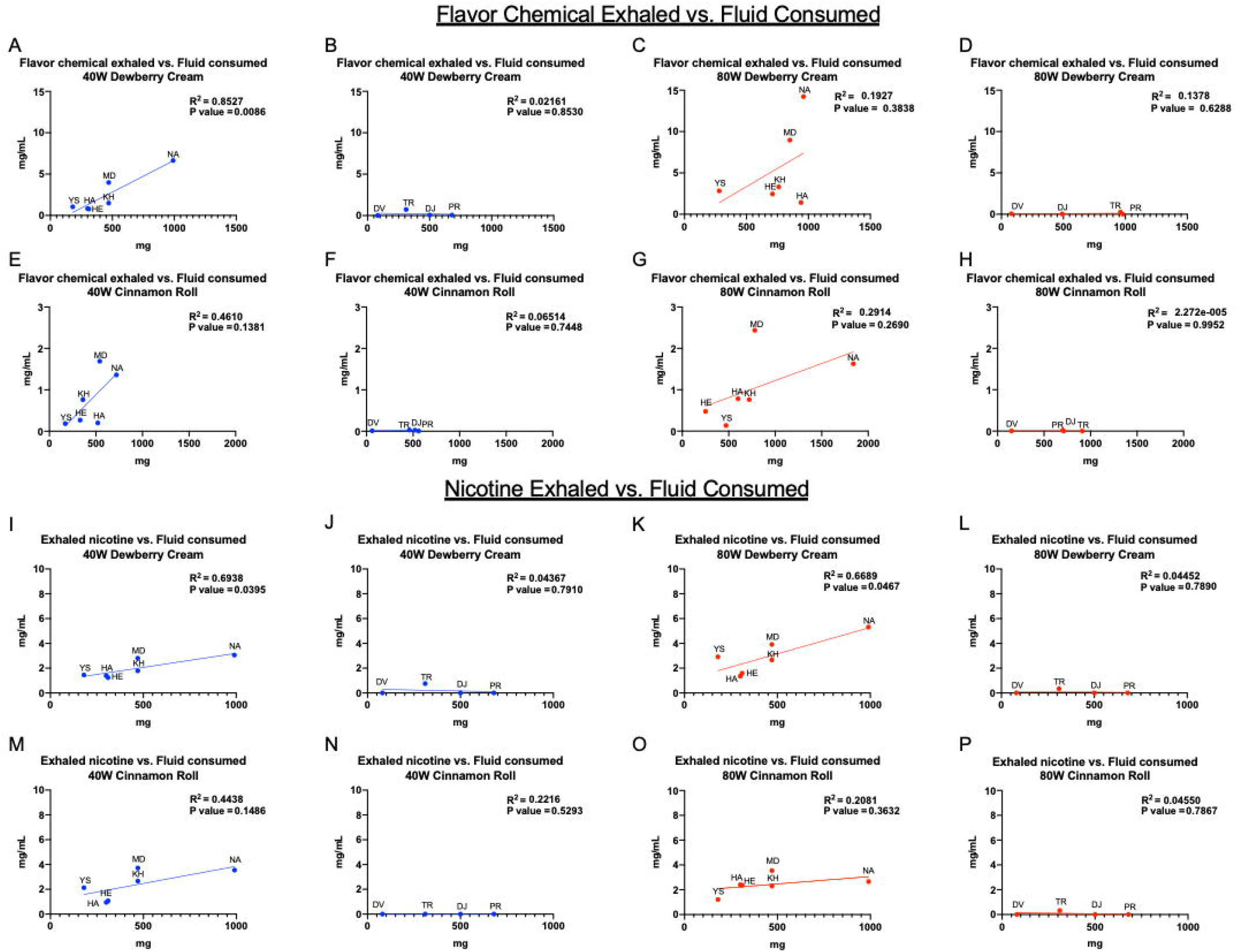
Participant’s Topography. Relationship between the refill fluid consumed and the flavor chemicals exhaled for both EC settings and refill fluids (A-H). Relationship between fluid consumed and nicotine exhaled for both EC settings and refill fluids (I-P). Mouth inhalers are shown in A, C, E, G, I, K, M, O and lung inhalers are shown in B, D, F, H, J, L, N, and P.

#### v. Fluid Consumed Vs. Nicotine Exhaled

Exhaled nicotine was quantified and compared to the amount of refill fluid consumed (Figure 4I-P). For the lung inhalers, there was no correlation between nicotine exhaled and the amount of fluid consumed (Fig. 4J, L, N, P). For the mouth inhalers, there was a significant correlation for the Dewberry Cream refill fluid and nicotine exhaled at both 40 (Fig. 4I) and 80 (Fig. 4K) watts, while there was no correlation for Cinnamon Roll at either wattage (4M and 4O).

#### vi. Percent Retention and Contribution of Exhale to ECEAR

We computed retention and ECEAR for each of the topographies. The percent retention was calculated and averaged for all participants (Figure 5). Lung inhalers had ~100% retention for flavor chemicals and nicotine for each setting/topography. Mouth inhalers retained variable percentages of specific flavor chemicals and nicotine. For mouth inhalers, cinnamaldehyde was retained better than nicotine and other flavor chemicals (Fig. 5G), and nicotine (Fig. 5E and 5F) was retained better than maltol, ethyl maltol, vanillin and ethyl vanillin (Fig. 5A-D).

**Figure 5:**
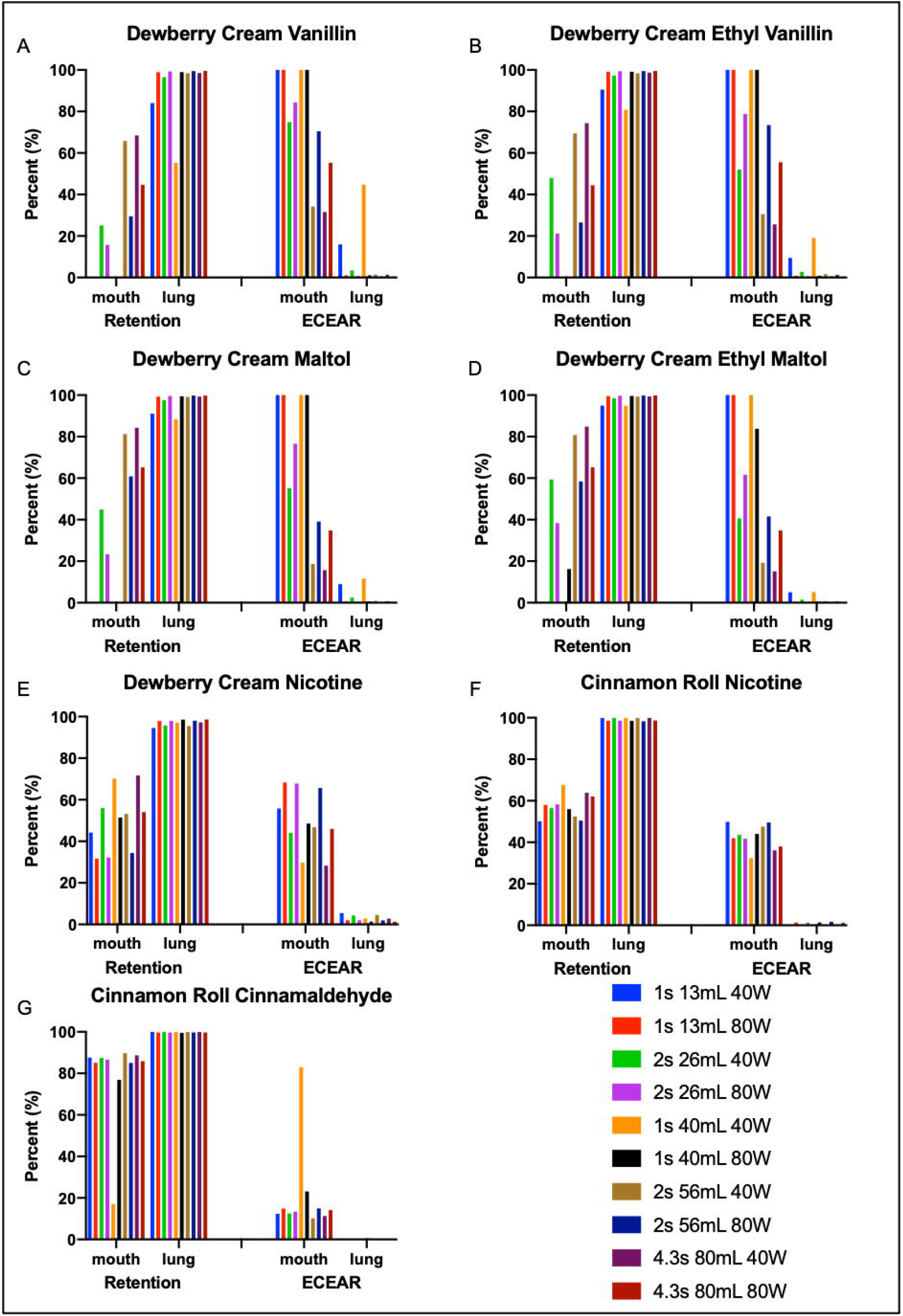
Retention and contribution to ECEAR of major flavor chemicals by participants under several EC settings and conditions. Each participant’s exhaled results were averaged for each topography to determine possible retention. Contribution to ECEAR was then calculated and averaged based on the amount retained. The y axis shows the percent retention or percent ECEAR while the x axis shows the participants averaged and separated by method of inhalation. EC settings, puff duration and puff volume are color coated.

The percent of inhaled aerosol that was exhaled and could contribute to ECEAR is also shown in Figure 5A-G. Lung inhalers did not contribute to ECEAR (Fig. 5A-G). However, mouth inhalers did contribute to ECEAR, and their contribution depended on flavor chemicals. Vanillin, ethyl vanillin, maltol, and ethyl maltol (Fig. 5A-D) contributed more to ECEAR than nicotine or cinnamaldehyde by mouth inhalers (Fig. 5E-G). There was very little contribution of cinnamaldehyde to ECEAR by mouth inhalers (Fig. 5G). The average nicotine contribution to ECEAR by mouth inhalers was 50% for “Dewberry Cream” and 42.5% for “Cinnamon Roll” (Fig. 5E, F).

#### vii. Concentrations of Flavor Chemical and Nicotine in Exhale

The concentrations of specific flavor chemicals and nicotine in the exhale of the mouth and lung inhalers is shown in Figure 6. In most cases, exhaled concentrations were higher when vaping was done at 40 W. Mouth inhalers exhaled nicotine and flavor chemicals at concentrations > 1 mg/mL (Fig. 6A-L), while concentrations for lung inhalers (Fig. 6M-T) were < 1mg/mL and were often not detectable.

**Figure 6:**
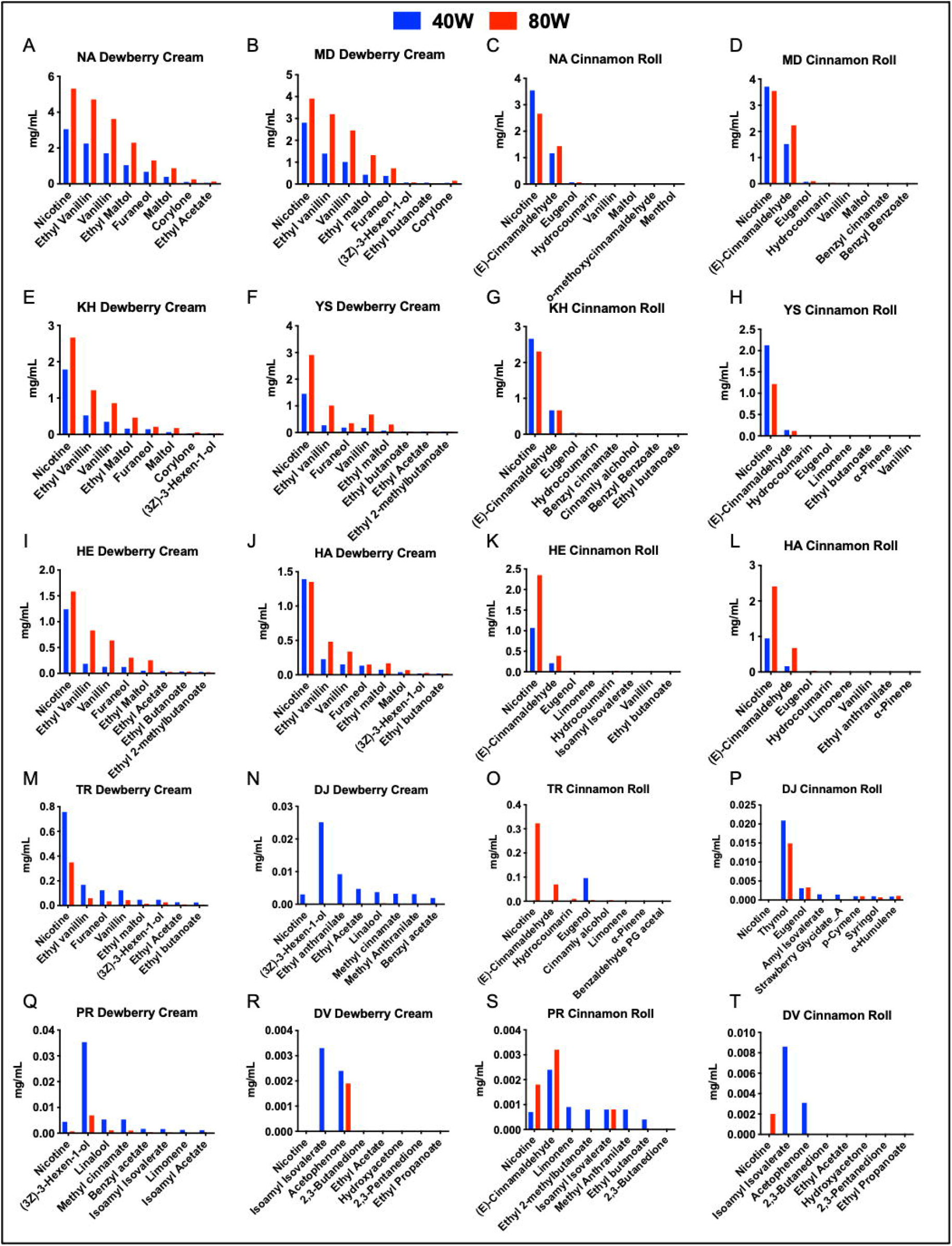
The concentration of major flavor chemicals emitted by each participant. 40-watt setting is shown in blue and 80-watt setting is shown in red.

## 4. DISCUSSION

To the best of our knowledge, this is the first study to trace the movement of flavor chemicals/nicotine from refill fluids to exhaled aerosol. The EC settings and flavor chemicals in each refill fluid effected transfer efficiency and chemical retention. Participants either exhaled little or no nicotine/flavor chemicals or they exhaled up to half of what was found in the refill fluid. We interpret this to mean that the former group inhaled aerosol into their lungs where chemicals were efficiently absorbed (lung inhalers), while the latter group kept much of the aerosol in their mouths, then exhaled aerosol only partially depleted of chemicals (mouth inhalers). This distinction is important since chemical exposure varied considerably between the two types of inhalers and only the mouth inhalers contributed nicotine and flavor chemicals to ECEAR.

The flavor chemicals in “Dewberry Cream” were similar to those reported previously (Hua et al., 2019), with some bottle-to-bottle variation in total flavor chemical concentration (24, 25 and 28 mg/mL). In contrast, there was about a 5-fold difference in cinnamaldehyde concentration in bottles of “Cinnamon Roll” (#537 = 13.4 mg/ml and #539 = 61.4 mg/ml) purchased at different times in a local vape shop, where the compounding was not precisely controlled. Maltol, ethyl maltol, vanillin, and ethyl vanillin were detected in high concentrations in “Dewberry Cream” and are among the most potent flavor chemicals when tested *in vitro* with mouse neural stem cells and BEAS-2B cells in the MTT assay (Hua et al., 2019). Cinnamaldehyde, while present in Cinnamon Roll, was low in concentration compared to other cinnamon flavored products we have examined (Behar et al., 2016).

Transfer efficiency of flavor chemicals and nicotine from machine-vaped refill fluid to aerosols depended on the properties of the chemicals, EC wattage, the pump head, puff duration and puff volume. Maltol, ethyl maltol, vanillin, and ethyl vanillin had similar patterns of transfer efficiency, which increased as puff volume, duration, and wattage increased. Nicotine transferred well and was not affected by these factors. Cinnamaldehyde and eugenol were similar to nicotine when the standard pump head was used. Of the chemicals tested, nicotine had the highest vapor pressure and hence lowest intermolecular forces (Table 2), which likely contributed to its high transfer efficiency. Eugenol and cinnamaldehyde had slightly lower vapor pressures, which may explain their efficient transfer with the standard pump head. However, like other flavor chemicals, eugenol and cinnamaldehyde did not transfer well with the high-performance pump head, probably due to the mechanics of the pump. For those chemicals with low vapor pressures (maltol, ethyl maltol, vanillin, and ethyl vanillin), transfer efficiency was also likely affected by the heat generated in the atomizers. Efficiency increased when puff duration and wattage increased, both factors which increase heat. These results fall in line with a study showing an increase in voltage from 3 to 6V increased aerosol generation across refill fluids and cartomizers (Havel et al., 2017). Although we tested only one brand of EC, transfer efficiencies would likely also be affected by EC brand.

**Table 2:**
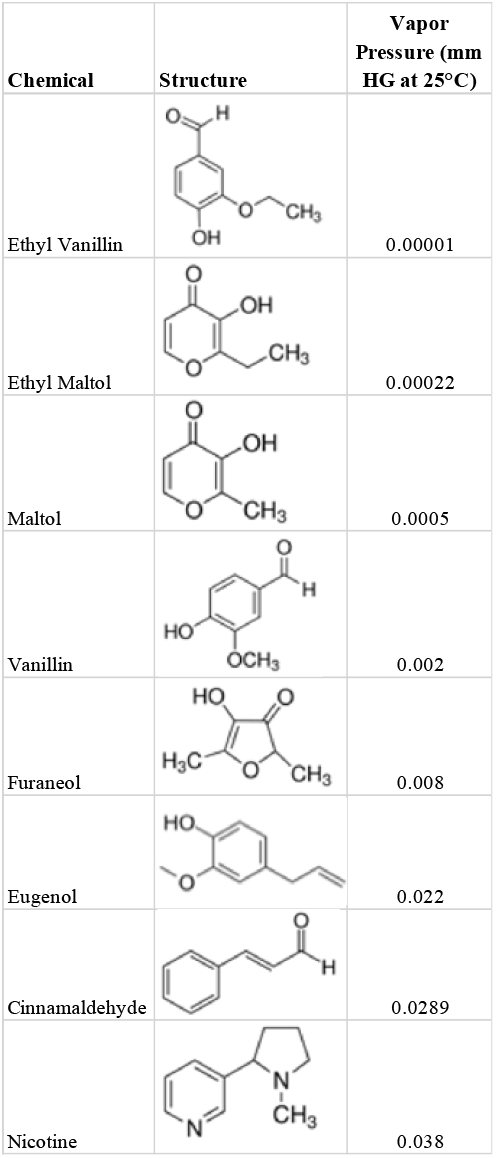
Vapor pressures and structures of flavor chemicals and nicotine.

| Chemical | Structure | Vapor Pressure (mm HG at 25°C) |
| --- | --- | --- |
| Ethyl Vanillin |  | 0.00001 |
| Ethyl Maltol |  | 0.00022 |
| Maltol |  | 0.0005 |
| Vanillin |  | 0.002 |
| Furaneol |  | 0.008 |
| Eugenol |  | 0.022 |
| Cinnamaldehyde |  | 0.0289 |
| Nicotine |  | 0.038 |

EC puffing topography varied among participants but was usually similar between trials for each individual, in agreement with Behar et al. 2015 who showed that each participant had their own “fingerprint” that defined their puffing topography (Behar et al., 2015). Our participants had similar patterns of puff volume and exhale irrespective of the wattage or refill fluid they were using. In a preliminary *ad libitum* study, users had an average of 3.5 ± 1.4 seconds puff duration (St Helen et al., 2016), while another study evaluated YouTube videos for an average of 4.3 seconds puff duration (Hua et al., 2013). In our study, the average puff duration (1.4 ± 0.27 seconds) could be related to the younger age of our participants and/or their lack of cigarette smoking experience.

The concentration of exhaled flavor chemicals increased when the ECs were operated at a higher wattage. Nicotine exhale also varied with the wattage and refill fluid consumed. Based on the exhale data, there were two categories of vapers – those who exhaled some flavor chemicals and those who exhaled little or no flavor chemicals. It has been suggested that naïve vapers using first generation ‘cig-a-like’ ECs had buccal rather than pulmonary absorption (Bullen et al., 2010; Vansickel et al., 2012). By quantifying the exhale of the participants, we were able to distinguish mouth vs. lung inhalation. Our participants were young (average age 21), and only one participant reported the use of tobacco cigarettes once a month. Therefore, it is possible that the “mouth inhalers” have not yet learned how to inhale into their lungs for maximum nicotine retention or they intentionally chose not to do this as they engage in vaping as a social activity.

We modeled chemical retention for 10 topographies (Fig. 5) and found that retention varied among chemicals and between user topographies (i.e., lung vs mouth inhalers). Cinnamaldehyde was retained better than other flavor chemicals by the “mouth inhalers”, suggesting that it is more soluble and/or reactive than the other aldehydes (e.g., vanillin or ethyl vanillin). This is concerning because cinnamaldehyde induces loss of ciliary motility and impairs mucociliary transport leading to respiratory infections (Clapp et al., 2019). Cinnamon-flavored refill fluids were also the most toxic of 36 refill fluids screened in vitro with three different cell types in the MTT assay (Bahl et al., 2012), and some cinnamon flavored products have very high concentrations of cinnamaldehyde, up to 343 mg/ml (Omaiye et al., 2019). It may be difficult for users to avoid exposure to cinnamaldehyde, as it has been reported in refill fluids that do not indicate a cinnamon flavor, such as Black Cherry or Caramel (Behar et al., 2016).

The retention of flavor chemicals and nicotine was about 100% for all “lung inhalers”, while retention for “mouth inhalers” was variable, but never 100%. In fact, nicotine was better retained than all flavor chemicals except cinnamaldehyde. These data that add to the information needed to evaluate human exposure to EC aerosols. The amount and rate of nicotine delivered may depend on the user topography, such as puff duration, or the nicotine concentration or the flavor (Dawkins and Corcoran, 2014; Hiler et al., 2017; Hejek et al., 2017; Helen et al., 2018; Voos et al., 2019). The retention of nicotine may be influenced by various factors, such as the pH of refill fluids or protonation by benzoic acid (Helen et al., 2018; Duell et al., 2018; Pankow et al., 2017), which is particularly relevant to pod-style products that contain acids and high nicotine concentrations.

Exhaled aerosol settles on surfaces forming ECEAR, which can remain for months (Khachatoorian et al., 2018). In previous studies, ECEAR had nicotine concentrations ranging from 0.03 to 0.949 μg/cm^2^ depending on the surface (Marcham et al., 2019), while an EC user’s home had 7.7±17.2 μg/m^2^ (Bush and Goniewicz, 2015). However, our previous study showed nicotine accumulated to a concentration of 108 mg/m^2^ after 1 month inside a vape shop and up to 1,181 μg/m^2^ inside a living room field site after 3 months (Khachatoorian et al., 2019). The exhaled flavor chemicals and nicotine in ECEAR are mostly likely contributed by mouth inhalers. Nevertheless, lung inhalers do exhale a visible puff of aerosol, which may contain mainly solvents. Other chemicals that were not measured in this study that could contribute to ECEAR include solvents, metals, and reaction products, such as formaldehyde, and acetaldehyde (Son et al., 2020; Li et al., 2020; Geiss et al., 2015).

While our focus was on ECEAR, the suspended exhale from EC users could also cause passive secondhand exposure to non-vapors. This idea is supported by studies in which non-vaping participants who were exposed to secondhand EC aerosols had alterations in respiratory mechanics and increases in salivary and urinary cotinine, urinary trans-3’-hydroxycotinine, and acrolein metabolites (Johnson et al., 2019; Tzortzi et al., 2018).

In conclusion, this is the first study to quantify flavor chemicals and nicotine in refill fluids, aerosols, and EC users’ exhale and then deduce their retention and contribution to ECEAR. The transfer of flavor chemicals with low vapor pressures to aerosols was dependent on puff duration, puff volume, user topography, pump head, and EC wattage, while nicotine transfer was not significantly affected by these factors. Analysis of exhaled chemicals enabled identification of mouth and lung inhalers. Mouth inhalers exhaled chemicals and contributed to ECEAR, while lung inhalers retained almost all the inhaled flavor chemicals and nicotine. Since the retention of toxic chemicals is higher in lung inhalers, harm reduction could be achieved if lung inhalers switched to mouth inhalation; however, this would increase the concentration of chemicals in ECEAR, which may affect those who are passively exposed to EC chemicals. These data contribute to our understanding of EC chemical transfer, retention, and contribution to ECEAR and are important to inform EC users, the public, and government agencies of potential exposures to chemical produced by ECs.

### LIMITATIONS OF THE STUDY

Our study is based on a relatively small sample size comprised of predominantly young

Asian males. Future studies could be expanded to include a more ethnically diverse population of EC users and more females. While our data are based on a single brand of EC, numerous brands spanning four generations are available, and should be evaluated in the future to determine how results are affected by brand. The introduction of pod-style ECs and loopholes in the flavor ban have led to the increased use of disposable pod-style ECs with many flavors and higher nicotine concentrations than were used in our study (US Dept. of Health & Human Services). Pod based products would be particularly interesting to examine in the future since these advanced devices can deliver higher nicotine concentrations to EC users (Yingst et al., 2019a;Yingst et al., 2019b).

## Resource Availability

- **Lead Contact:** Further information and requests for resources and reagents should be directed to and will be fulfilled by the Lead Contact, Prue Talbot.
- **Materials Available:** All relevant data are included in the manuscript and supplement.
- **Data and Code Availability:** All raw data are available upon request to the Lead Contact.

## Acknowledgments

We thank Riaz Golshan and Omeka Ikegbu for generation of some of the aerosols.

## Contributors

Conception and design: CK and PT. Participant recruiting: CK and PT. Sample preparation and data collection: CK. Chemical analysis: WL, KJM, and JP. Data interpretation: CK, WL, KJM, JP, and PT. Data analysis and writing of the manuscript: CK and PT. Editing the manuscript: CK, WL, KJM, JP, and PT.

## Supplementary Figures

**Figure S1:**
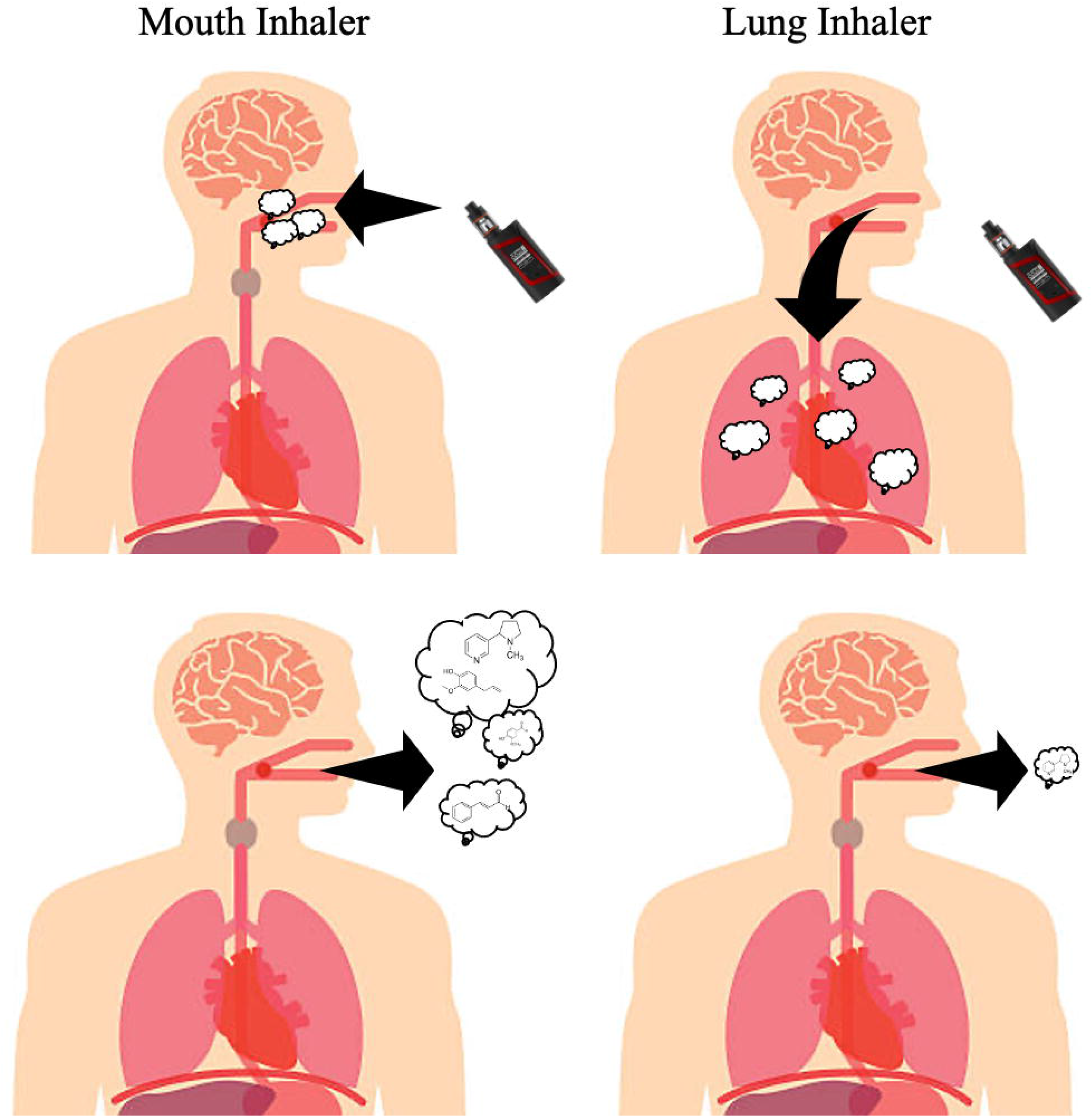
Heatmaps of each participant’s exhale with “Dewberry Cream” and “Cinnamon roll” used at 40 and 80 watts.

**Table S1:** Participant’s control exhale (no EC use).

