## Supplementary Data for "Tracking the Movement of Electronic Cigarette Flavor Chemicals and Nicotine from Refill Fluids to Aerosol, Lungs and Exhale"

Supplementary Table 1: Control Exhale Data (no EC use)

|  | DV CN Exhale | YS CN Exhale | HA CN Exhale | HE CN Exhale | KH CN Exhale | TR CN Exhale | MD CN Exhale | DJ CN Exhale | PR CN Exhale | NA CN Exhale |
| --- | --- | --- | --- | --- | --- | --- | --- | --- | --- | --- |
| **Compound Name** |  |  |  |  |  |  |  |  |  |  |
| 2,3-Butanedione | 0 | 0 | 0 | 0 | 0 | 0 | 0 | 0 | 0 | 0 |
| Ethyl Acetate | 0 | 0 | 0 | 0 | 0 | 0 | 0 | 0 | 0 | 0 |
| Hydroxyacetone | 0 | 0 | 0 | 0 | 0 | 0 | 0 | 0 | 0 | 0 |
| 2,3-Pentanedione | 0 | 0 | 0 | 0 | 0 | 0 | 0 | 0 | 0 | 0 |
| Ethyl Propanoate | 0 | 0 | 0 | 0 | 0 | 0 | 0 | 0 | 0 | 0 |
| Acetoin | 0 | 0 | 0 | 0 | 0 | 0 | 0 | 0 | 0 | 0 |
| Ethyl Isobutyrate | 0 | 0 | 0 | 0 | 0 | 0 | 0 | 0 | 0 | 0 |
| Isopentyl Alcohol | 0 | 0 | 0 | 0 | 0 | 0 | 0 | 0 | 0 | 0 |
| Isobutyl Acetate | 0 | 0 | 0 | 0 | 0 | 0 | 0 | 0 | 0 | 0 |
| Methyl 2-methylbutyrate | 0 | 0 | 0 | 0 | 0 | 0 | 0 | 0 | 0 | 0 |
| 1-Pentanol | 0 | 0 | 0 | 0 | 0 | 0 | 0 | 0 | 0 | 0 |
| 2,3-Hexanedione | 0 | 0 | 0 | 0 | 0 | 0 | 0 | 0 | 0 | 0 |
| Ethyl butanoate | 0 | 0 | 0 | 0 | 0 | 0 | 0 | 0 | 0 | 0 |
| Hexanal | 0 | 0 | 0 | 0 | 0 | 0 | 0 | 0 | 0 | 0 |
| Butyl Acetate | 0 | 0 | 0 | 0 | 0 | 0 | 0 | 0 | 0 | 0 |
| Ethyl lactate | 0 | 0 | 0 | 0 | 0 | 0 | 0 | 0 | 0 | 0 |
| Ethyl 2-methylbutanoate | 0 | 0 | 0 | 0 | 0 | 0 | 0 | 0 | 0 | 0 |
| Ethyl Isovalerate | 0 | 0 | 0 | 0 | 0 | 0 | 0 | 0 | 0 | 0 |
| Furfural | 0 | 0 | 0 | 0 | 0 | 0 | 0 | 0 | 0 | 0 |
| Isoamyl Acetate | 0 | 0 | 0 | 0 | 0 | 0 | 0 | 0 | 0 | 0 |
| 2-Methylbutyl Acetate | 0 | 0 | 0 | 0 | 0 | 0 | 0 | 0 | 0 | 0 |
| (3Z)-3-Hexen-1-ol | 0 | 0 | 0 | 0 | 0 | 0 | 0 | 0 | 0 | 0 |
| 2-Hexen-1-ol, (E)- | 0 | 0 | 0 | 0 | 0 | 0 | 0 | 0 | 0 | 0 |
| 1-Hexanol | 0 | 0 | 0 | 0 | 0 | 0 | 0 | 0 | 0 | 0 |
| Furfuryl alcohol | 0 | 0 | 0 | 0 | 0 | 0 | 0 | 0 | 0 | 0 |
| Amyl Acetate | 0 | 0 | 0 | 0 | 0 | 0 | 0 | 0 | 0 | 0 |
| 2,5-dimethylpyrazine | 0 | 0 | 0 | 0 | 0 | 0 | 0 | 0 | 0 | 0 |
| (3Z)-3-Hexenyl Formate | 0 | 0 | 0 | 0 | 0 | 0 | 0 | 0 | 0 | 0 |
| α-Pinene | 0 | 0 | 0 | 0 | 0 | 0 | 0 | 0 | 0 | 0 |
| 2,3-dimethylpyrazine | 0 | 0 | 0 | 0 | 0 | 0 | 0 | 0 | 0 | 0 |
| Ethyl 3-hydroxybutyrate | 0 | 0 | 0 | 0 | 0 | 0 | 0 | 0 | 0 | 0 |
| Isoamyl Propionate | 0 | 0 | 0 | 0 | 0 | 0 | 0 | 0 | 0 | 0 |
| 2-Methoxy-3-methylpyrazine | 0 | 0 | 0 | 0 | 0 | 0 | 0 | 0 | 0 | 0 |
| β-Myrcene | 0 | 0 | 0 | 0 | 0 | 0 | 0 | 0 | 0 | 0 |
| β-Pinene | 0 | 0 | 0 | 0 | 0 | 0 | 0 | 0 | 0 | 0 |
| Butyl butyrate | 0 | 0 | 0 | 0 | 0 | 0 | 0 | 0 | 0 | 0 |
| Ethyl hexanoate | 0 | 0 | 0 | 0 | 0 | 0 | 0 | 0 | 0 | 0 |
| Benzaldehyde | 0 | 0 | 0 | 0 | 0 | 0 | 0 | 0 | 0 | 0 |
| Pentyl Propanoate | 0 | 0 | 0 | 0 | 0 | 0 | 0 | 0 | 0 | 0 |
| 6-Methyl-5-heptene-2-one | 0 | 0 | 0 | 0 | 0 | 0 | 0 | 0 | 0 | 0 |
| 3-Hexen-1-ol, acetate, (Z)- | 0 | 0 | 0 | 0 | 0 | 0 | 0 | 0 | 0 | 0 |
| 2,3,5-Trimethylpyrazine | 0 | 0 | 0 | 0 | 0 | 0 | 0 | 0 | 0 | 0 |
| Hexyl acetate | 0 | 0 | 0 | 0 | 0 | 0 | 0 | 0 | 0 | 0 |
| 2-ethyl-3-methylpyrazine | 0 | 0 | 0 | 0 | 0 | 0 | 0 | 0 | 0 | 0 |
| 1,4-Cineol | 0 | 0 | 0 | 0 | 0 | 0 | 0 | 0 | 0 | 0 |
| Limonene | 0 | 0 | 0 | 0 | 0 | 0 | 0 | 0 | 0 | 0 |
| p-Cymene | 0 | 0 | 0 | 0 | 0 | 0 | 0 | 0 | 0 | 0 |
| γ-Pentalactone | 0 | 0 | 0 | 0 | 0 | 0 | 0 | 0 | 0 | 0 |
| Eucalyptol | 0 | 0 | 0 | 0 | 0 | 0 | 0 | 0 | 0 | 0 |
| γ-Terpinene | 0 | 0 | 0 | 0 | 0 | 0 | 0 | 0 | 0 | 0 |
| Dimethyl butanedioate | 0 | 0 | 0 | 0 | 0 | 0 | 0 | 0 | 0 | 0 |
| Acetylpyrazine | 0 | 0 | 0 | 0 | 0 | 0 | 0 | 0 | 0 | 0 |
| Isoamyl Butyrate | 0 | 0 | 0 | 0 | 0 | 0 | 0 | 0 | 0 | 0 |
| Corylone | 0 | 0 | 0 | 0 | 0 | 0 | 0 | 0 | 0 | 0 |
| Ally hexanoate | 0 | 0 | 0 | 0 | 0 | 0 | 0 | 0 | 0 | 0 |
| Benzeneacetaldehyde | 0 | 0 | 0 | 0 | 0 | 0 | 0 | 0 | 0 | 0 |
| Benzyl Alcohol | 0 | 0 | 0 | 0 | 0 | 0 | 0 | 0 | 0 | 0 |
| Amyl Butyrate | 0 | 0 | 0 | 0 | 0 | 0 | 0 | 0 | 0 | 0 |
| 2,3,5,6-Tetramethylpyrazine | 0 | 0 | 0 | 0 | 0 | 0 | 0 | 0 | 0 | 0 |
| Ethyl Heptanoate | 0 | 0 | 0 | 0 | 0 | 0 | 0 | 0 | 0 | 0 |
| cis-Linalool oxide | 0 | 0 | 0 | 0 | 0 | 0 | 0 | 0 | 0 | 0 |
| Furaneol | 0 | 0 | 0 | 0 | 0 | 0 | 0 | 0 | 0 | 0 |
| Isoamyl Isovalerate | 0 | 0 | 0 | 0 | 0 | 0 | 0 | 0 | 0 | 0 |
| Amyl Isovalerate | 0 | 0 | 0 | 0 | 0 | 0 | 0 | 0 | 0 | 0 |
| trans-Linalool Oxide | 0 | 0 | 0 | 0 | 0 | 0 | 0 | 0 | 0 | 0 |
| 2-Nonanone | 0 | 0 | 0 | 0 | 0 | 0 | 0 | 0 | 0 | 0 |
| Acetophenone | 0 | 0 | 0 | 0 | 0 | 0.3 | 0 | 0 | 0 | 0.2 |
| Linalool | 0 | 0 | 0 | 0 | 0 | 0 | 0 | 0 | 0 | 0 |
| 2-Acetylpyrrole | 0 | 0 | 0 | 0 | 0 | 0 | 0 | 0 | 0 | 0 |
| p-Tolualdehyde | 0 | 0 | 0 | 0 | 0 | 0 | 0 | 0 | 0 | 0 |
| Guaiacol (2-methoxyphenol) | 0 | 0 | 0 | 0 | 0 | 0 | 0 | 0 | 0 | 0 |
| 2-Methylbenzofuran | 0 | 0 | 0 | 0 | 0 | 0 | 0 | 0 | 0 | 0 |
| 1,2-Dihydrolinalool | 0 | 0 | 0 | 0 | 0 | 0 | 0 | 0 | 0 | 0 |
| cis-Limonene oxide | 0 | 0 | 0 | 0 | 0 | 0 | 0 | 0 | 0 | 0 |
| Fenchol | 0 | 0 | 0 | 0 | 0 | 0 | 0 | 0 | 0 | 0 |
| trans-Limonene oxide | 0 | 0 | 0 | 0 | 0 | 0 | 0 | 0 | 0 | 0 |
| Maltol | 0 | 0 | 0 | 0 | 0 | 0 | 0 | 0 | 0 | 0 |
| Phenethyl alcohol | 0 | 0 | 0 | 0 | 0 | 0 | 0 | 0 | 0 | 0 |
| β-Citronellal | 0 | 0 | 0 | 0 | 0 | 0 | 0 | 0 | 0 | 0 |
| Isopulegol | 0 | 0 | 0 | 0 | 0 | 0 | 0 | 0 | 0 | 0 |
| 4-methylbenzyl alcohol | 0 | 0 | 0 | 0 | 0 | 0 | 0 | 0 | 0 | 0 |
| Benzyl acetate | 0 | 0 | 0 | 0 | 0 | 0 | 0 | 0 | 0 | 0 |
| Ethyl octanoate | 0 | 0 | 0 | 0 | 0 | 0 | 0 | 0 | 0 | 0 |
| 2-Hydroxy-3,5,5-trimethyl-cyclohex-2-en | 0 | 0 | 0 | 0 | 0 | 0 | 0 | 0 | 0 | 0 |
| p-Menthone | 0 | 0 | 0 | 0 | 0 | 0 | 0 | 0 | 0 | 0 |
| Ethyl Benzoate | 0 | 0 | 0 | 0 | 0 | 0 | 0 | 0 | 0 | 0 |
| Neomenthol | 0 | 0 | 0 | 0 | 0 | 0 | 0 | 0 | 0 | 0 |
| Methyl phenylacetate | 0 | 0 | 0 | 0 | 0 | 0 | 0 | 0 | 0 | 0 |
| 4-Terpineol | 0 | 0 | 0 | 0 | 0 | 0 | 0 | 0 | 0 | 0 |
| Menthol | 0 | 0 | 0 | 0 | 0 | 1.4 | 0 | 0 | 0 | 0 |
| Styralyl Acetate | 0 | 0 | 0 | 0 | 0 | 0 | 0 | 0 | 0 | 0 |
| Benzylacetaldehyde | 0 | 0 | 0 | 0 | 0 | 0 | 0 | 0 | 0 | 0 |
| Methyl 2-Octynate | 0 | 0 | 0 | 0 | 0 | 0 | 0 | 0 | 0 | 0 |
| Estragole (4-allyanisole) | 0 | 0 | 0 | 0 | 0 | 0 | 0 | 0 | 0 | 0 |
| 3'-Methylacetophenone | 0 | 0 | 0 | 0 | 0 | 0 | 0 | 0 | 0 | 0 |
| α-Terpineol | 0 | 0 | 0 | 0 | 0 | 0 | 0 | 0 | 0 | 0 |
| Hexyl 2-Methylbutyrate | 0 | 0 | 0 | 0 | 0 | 0 | 0 | 0 | 0 | 0 |
| Methyl salicylate | 0 | 0 | 0 | 0 | 0 | 0 | 0 | 0 | 0 | 0 |
| γ-Heptalactone | 0 | 0 | 0 | 0 | 0 | 0 | 0 | 0 | 0 | 0 |
| Linaly acetate | 0 | 0 | 0 | 0 | 0 | 0 | 0 | 0 | 0 | 0 |
| Ethyl Maltol | 0 | 0 | 0 | 0 | 0 | 0 | 0 | 0 | 0 | 0 |
| Benzeneacetic acid, ethyl ester | 0 | 0 | 0 | 0 | 0 | 0 | 0 | 0 | 0 | 0 |
| trans-Geraniol | 0 | 0 | 0 | 0 | 0 | 0 | 0 | 0 | 0 | 0 |
| Benzyl Propionate | 0 | 0 | 0 | 0 | 0 | 0 | 0 | 0 | 0 | 0 |
| Pulegone | 0 | 0 | 0 | 0 | 0 | 0 | 0 | 0 | 0 | 0 |
| Ethyl nonanoate | 0 | 0 | 0 | 0 | 0 | 0 | 0 | 0 | 0 | 0 |
| Carvone | 0 | 0 | 0 | 0 | 0 | 0 | 0 | 0 | 0 | 0 |
| Menthyl Acetate | 0 | 0 | 0 | 0 | 0 | 0 | 0 | 0 | 0 | 0 |
| Benzaldehyde PG acetal | 0 | 0 | 0 | 0 | 0 | 0 | 0 | 0 | 0 | 0 |
| Citral | 0 | 0 | 0 | 0 | 0 | 0 | 0 | 0 | 0 | 0 |
| Ethyl salicylate | 0 | 0 | 0 | 0 | 0 | 0 | 0 | 0 | 0 | 0 |
| Piperitone | 0 | 0 | 0 | 0 | 0 | 0 | 0 | 0 | 0 | 0 |
| p-Anisaldehyde | 0 | 0 | 0 | 0 | 0 | 0 | 0 | 0 | 0 | 0 |
| Linaly propionate | 0 | 0 | 0 | 0 | 0 | 0 | 0 | 0 | 0 | 0 |
| Cinnamaldehyde, (E)- | 0 | 0 | 0 | 0 | 0 | 0 | 0 | 0 | 0 | 0 |
| Thymol | 0 | 0 | 0 | 0 | 0 | 0 | 0 | 0 | 0 | 0 |
| γ-Octalactone | 0 | 0 | 0 | 0 | 0 | 0 | 0 | 0 | 0 | 0 |
| Hemineurine | 0 | 0 | 0 | 0 | 0 | 0 | 0 | 0 | 0 | 0 |
| Nerol acetate | 0 | 0 | 0 | 0 | 0 | 0 | 0 | 0 | 0 | 0 |
| Butyl butyrolactate | 0 | 0 | 0 | 0 | 0 | 0 | 0 | 0 | 0 | 0 |
| Benzyl Butyrate | 0 | 0 | 0 | 0 | 0 | 0 | 0 | 0 | 0 | 0 |
| Cinnamly alchohol | 0 | 0 | 0 | 0 | 0 | 0 | 0 | 0 | 0 | 0 |
| Triacetin | 0 | 0 | 0 | 0 | 0 | 0 | 0 | 0 | 0 | 0 |
| Geraniol Acetate | 0 | 0 | 0 | 0 | 0 | 0 | 0 | 0 | 0 | 0 |
| Hexyl hexanoate | 0 | 0 | 0 | 0 | 0 | 0 | 0 | 0 | 0 | 0 |
| Ethyl decanoate | 0 | 0 | 0 | 0 | 0 | 0 | 0 | 0 | 0 | 0 |
| Nicotine | 0 | 0.2 | 0 | 0.7 | 0 | 1.6 | 0 | 0.7 | 0 | 0 |
| Eugenol | 0 | 0 | 0 | 0 | 0 | 0 | 0 | 0 | 0 | 0 |
| Syringol | 0 | 0 | 0 | 0 | 0 | 0 | 0 | 0 | 0 | 0 |
| Methyl Anthranilate | 0 | 0 | 0 | 0 | 0 | 0 | 0 | 0 | 0 | 0 |
| Piperonal | 0 | 0 | 0 | 0 | 0 | 0 | 0 | 0 | 0 | 0 |
| Eugenol methyl ether | 0 | 0 | 0 | 0 | 0 | 0 | 0 | 0 | 0 | 0 |
| Methyl cinnamate | 0 | 0 | 0 | 0 | 0 | 0 | 0 | 0 | 0 | 0 |
| α-Damascone | 0 | 0 | 0 | 0 | 0 | 0 | 0 | 0 | 0 | 0 |
| Ethyl Benzolyformate | 0 | 0 | 0 | 0 | 0 | 0 | 0 | 0 | 0 | 0 |
| Citronellyl Propionate | 0 | 0 | 0 | 0 | 0 | 0 | 0 | 0 | 0 | 0 |
| β-Caryophyllene | 0 | 0 | 0 | 0 | 0 | 0 | 0 | 0 | 0 | 0 |
| γ-Nonalactone | 0 | 0 | 0 | 0 | 0 | 0 | 0 | 0 | 0 | 0 |
| Methyl N-methylanthranilate | 0 | 0 | 0 | 0 | 0 | 0 | 0 | 0 | 0 | 0 |
| β-Damascone | 0 | 0 | 0 | 0 | 0 | 0 | 0 | 0 | 0 | 0 |
| Aromadendrene | 0 | 0 | 0 | 0 | 0 | 0 | 0 | 0 | 0 | 0 |
| α-Ionone | 0 | 0 | 0 | 0 | 0 | 0 | 0 | 0 | 0 | 0 |
| Ethyl anthranilate | 0 | 0 | 0 | 0 | 0 | 0 | 0 | 0 | 0 | 0 |
| Strawberry Glycidate_A | 0 | 0 | 0 | 0 | 0 | 0 | 0 | 0 | 0 | 0 |
| Cinnamyl Acetate | 0 | 0 | 0 | 0 | 0 | 0 | 0 | 0 | 0 | 0 |
| Hydrocoumarin | 0 | 0 | 0 | 0 | 0 | 0 | 0 | 0.6 | 0 | 0 |
| α-Caryophyllene (α-Humulene) | 0 | 0 | 0 | 0 | 0 | 0 | 0 | 0 | 0 | 0 |
| Vanillin | 0 | 0 | 0 | 0 | 0 | 0 | 0 | 0 | 0 | 0 |
| Ethyl Cinnamate | 0 | 0 | 0 | 0 | 0 | 0 | 0 | 0 | 0 | 0 |
| Benzyl dimethylcarbinyl butyrate | 0 | 0 | 0 | 0 | 0 | 0 | 0 | 0 | 0 | 0 |
| Isopentyl Phenylacetate | 0 | 0 | 0 | 0 | 0 | 0 | 0 | 0 | 0 | 0 |
| Isoeugenol methyl ether | 0 | 0 | 0 | 0 | 0 | 0 | 0 | 0 | 0 | 0 |
| (E)-β-Ionone | 0 | 0 | 0 | 0 | 0 | 0 | 0 | 0 | 0 | 0 |
| Ethyl Vanillin | 0 | 0 | 0 | 0 | 0 | 0 | 0 | 0 | 0 | 0 |
| Coumarin | 0 | 0 | 0 | 0 | 0 | 0 | 0 | 0 | 0 | 0 |
| Myristicin | 0 | 0 | 0 | 0 | 0 | 0 | 0 | 0 | 0 | 0 |
| γ-Decalactone | 0 | 0 | 0 | 0 | 0 | 0 | 0 | 0 | 0 | 0 |
| Acetyleugenol | 0 | 0 | 0 | 0 | 0 | 0 | 0 | 0 | 0 | 0 |
| Raspberry Ketone methyl ether | 0 | 0 | 0 | 0 | 0 | 0 | 0 | 0 | 0 | 0 |
| Hexyl octanoate | 0 | 0 | 0 | 0 | 0 | 0 | 0 | 0 | 0 | 0 |
| Strawberry Glycidate_B | 0 | 0 | 0 | 0 | 0 | 0 | 0 | 0 | 0 | 0 |
| Isosafroeugenol | 0 | 0 | 0 | 0 | 0 | 0 | 0 | 0 | 0 | 0 |
| Ethyl Laurate | 0 | 0 | 0 | 0 | 0 | 0 | 0 | 0 | 0 | 0 |
| δ-Decalactone | 0 | 0 | 0 | 0 | 0 | 0 | 0 | 0 | 0 | 0 |
| o-methoxycinnamaldehyde | 0 | 0 | 0 | 0 | 0 | 0 | 0 | 0 | 0 | 0 |
| δ-Undecalactone | 0 | 0 | 0 | 0 | 0 | 0 | 0 | 0 | 0 | 0 |
| Raspberry ketone | 0 | 0 | 0 | 0 | 0 | 0 | 0 | 0 | 0 | 0 |
| Coumarin, 6-methyl | 0 | 0 | 0 | 0 | 0 | 0 | 0 | 0 | 0 | 0 |
| Heliotropine proplyene gycol acetal | 0 | 0 | 0 | 0 | 0 | 0 | 0 | 0 | 0 | 0 |
| Benzyl ether | 0 | 0 | 0 | 0 | 0 | 0 | 0 | 0 | 0 | 0 |
| Benzophenone | 0 | 0 | 0 | 0 | 0 | 0 | 0 | 0 | 0 | 0 |
| Gingerone | 0 | 0 | 0 | 0 | 0 | 0 | 0 | 0 | 0 | 0 |
| δ-Dodecalactone | 0 | 0 | 0 | 0 | 0 | 0 | 0 | 0 | 0 | 0 |
| Benzyl Benzoate | 0 | 0 | 0 | 0 | 0 | 0 | 0 | 0 | 0 | 0 |
| Benzyl Benzeneacetate | 0 | 0 | 0 | 0 | 0 | 0 | 0 | 0 | 0 | 0 |
| Benzoin Ethyl Ether | 0 | 0 | 0 | 0 | 0 | 0 | 0 | 0 | 0 | 0 |
| Caffeine | 0 | 0 | 0 | 0 | 0 | 0 | 0 | 0 | 0 | 0 |
| Benzyl cinnamate | 0 | 0 | 0 | 0 | 0 | 0 | 0 | 0 | 0 | 0 |
| total µg | 0 | 0.2 | 0 | 0.7 | 0 | 3.3 | 0 | 1.3 | 0 | 0.2 |
| total mg | 0 | 0.0002 | 0 | 0.0007 | 0 | 0.0033 | 0 | 0.0013 | 0 | 0.0002 |

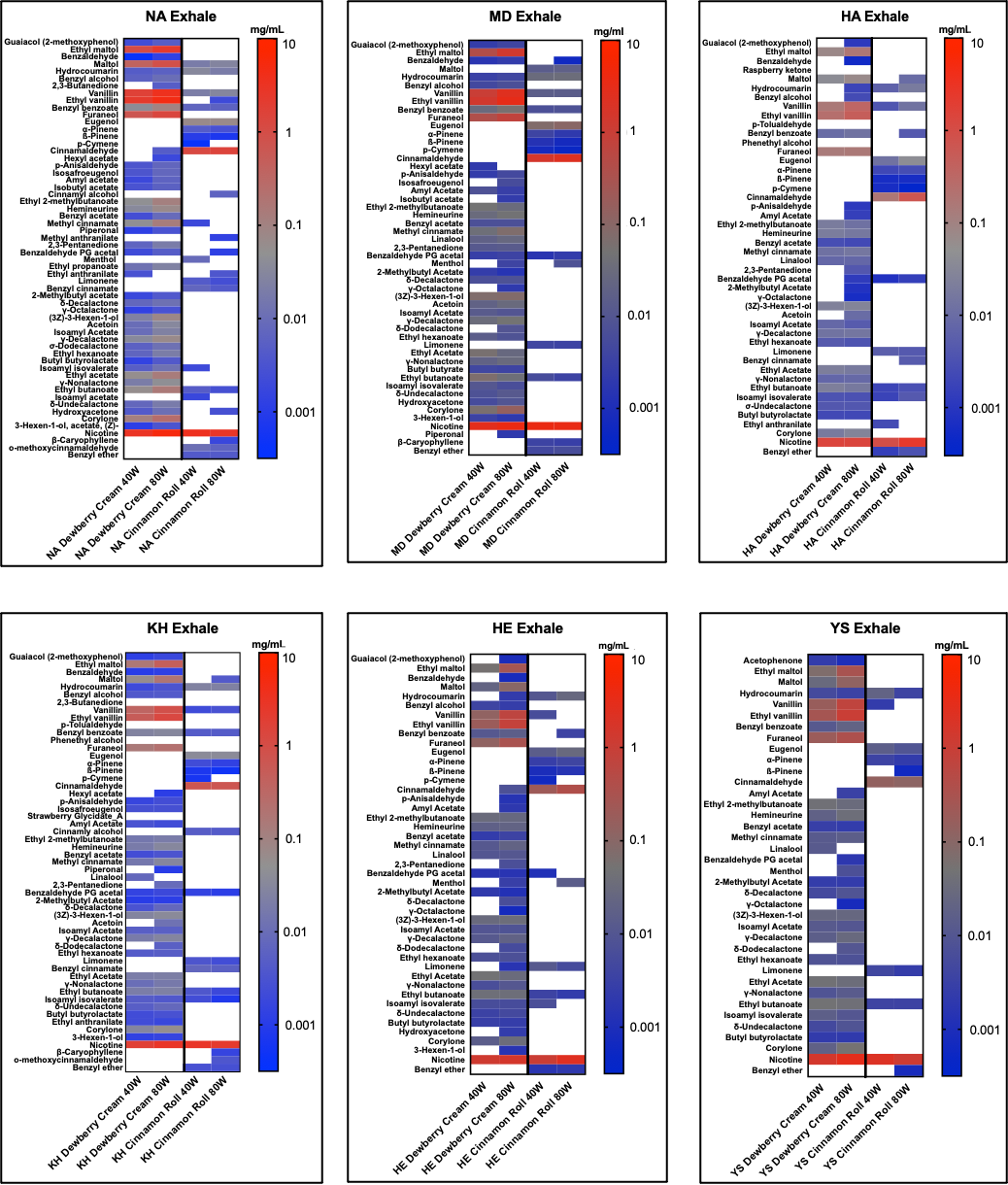
**Supplementary Figure 1:** Heatmaps of each participant’s exhale with “Dewberry Cream” #518 and “Cinnamon roll” #537 used at 40 and 80 watts.

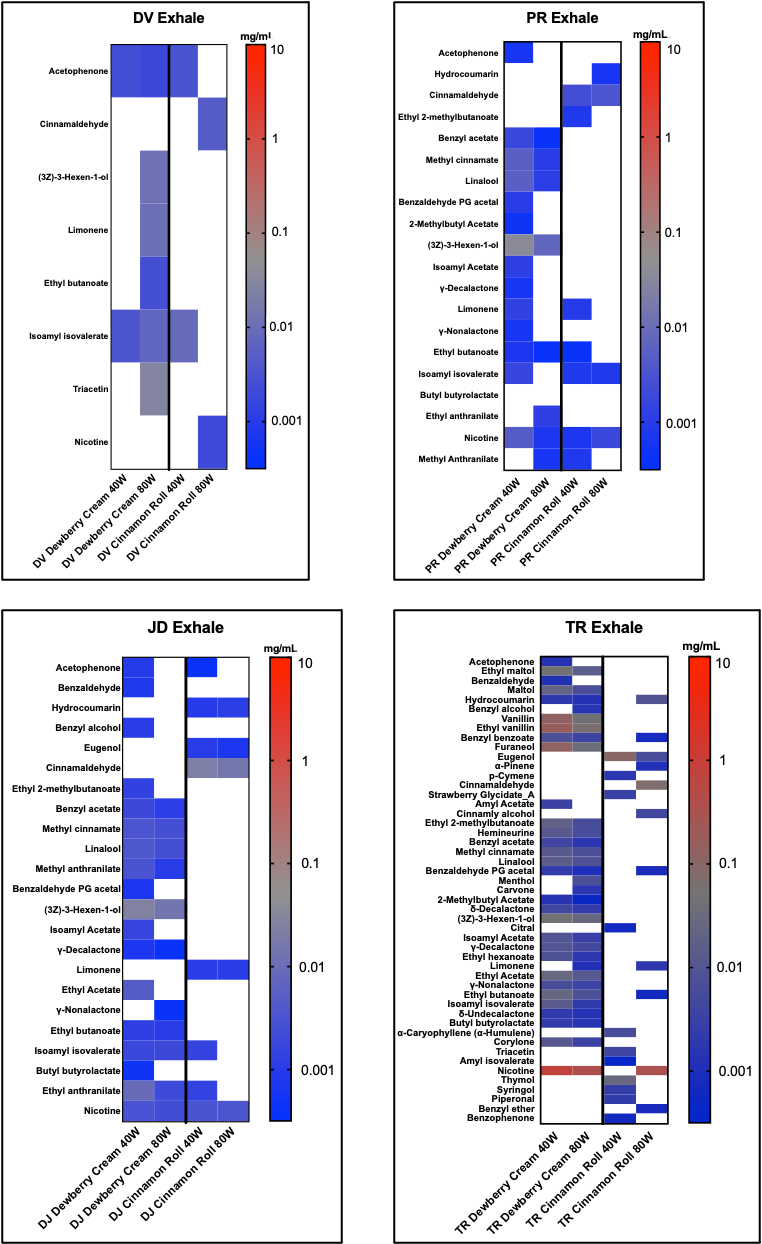
